# Estimating the temporal and spatial extent of gene flow among sympatric lizard populations (genus *Sceloporus*) in the southern Mexican highlands

**DOI:** 10.1101/008623

**Authors:** Jared A. Grummer, Martha L. Calderón-Espinosa, Adrián Nieto-Montes de Oca, Eric N. Smith, Fausto R. Méndez-de la Cruz, Adam D. Leaché

**Affiliations:** Department of Biology and Burke Museum of Natural History and Culture, University of Washington, Box 351800, Seattle, WA 98195-1800, USA; Instituto de Ciencias Naturales, Universidad Nacional de Colombia, Sede Bogotá Colombia; Museo de Zoología and Departmento de Biología Evolutiva, Facultad de Ciencias, Universidad Nacional Autónoma de México, Ciudad Universitaria, México 04510, Distrito Federal, México; Amphibian & Reptile Diversity Research Center, Department of Biology, University of Texas at Arlington, Arlington, TX 76019-0498; Laboratorio de Herpetología, Instituto de Biología, Universidad Nacional Autónoma de México, Ciudad Universitaria, México, D. F., C. P. 04510, México

**Keywords:** Mexico, mito-nuclear discordance, Bayesian phylogeography, hybridization, gene flow, coalescent simulations, species delimitation

## Abstract

Interspecific gene flow is pervasive throughout the tree of life. Although detecting gene flow between populations has been facilitated by new analytical approaches, determining the timing and geography of hybridization has remained difficult, particularly for historical gene flow. A geographically explicit phylogenetic approach is needed to determine the ancestral population overlap. In this study, we performed population genetic analyses, species delimitation, simulations, and a recently developed approach of species tree diffusion to infer the phylogeographic history, timing and geographic extent of gene flow in lizards of the *Sceloporus spinosus* group. The two species in this group, *S. spinosus* and *S. horridus*, are distributed in eastern and western portions of Mexico, respectively, but populations of these species are sympatric in the southern Mexican highlands. We generated data consisting of three mitochondrial genes and eight nuclear loci for 148 and 68 individuals, respectively. We delimited six lineages in this group, but found strong evidence of mito-nuclear discordance in sympatric populations of *S. spinosus* and *S. horridus* owing to mitochondrial introgression. We used coalescent simulations to differentiate ancestral gene flow from secondary contact, but found mixed support for these two models. Bayesian phylogeography indicated more than 60% range overlap between ancestral *S. spinosus* and *S. horridus* populations since the time of their divergence. Isolation-migration analyses, however, revealed near-zero levels of gene flow between these ancestral populations. Interpreting results from both simulations and empirical data indicate that despite a long history of sympatry among these two species, gene flow in this group has only recently occurred.

## Introduction

The topic of hybridization, or gene flow between evolutionary independent lineages, has captivated evolutionary biologists for nearly two centuries (Darwin 1859; Harrison 1993). Gene flow between species is common in nature with approximately 10% and 25% of animal and plant species known to hybridize, respectively (Mallet 2005). Although hybrid zones have been identified across a variety of organisms (Abbott et al 2013; Larson et al 2013), determining the temporal and geographic extent of hybridization has remained a difficult task (Hewitt 2001).

Analytical advancements in the field of phylogeography have enabled sophisticated model-testing approaches, including the ability to test demographic scenarios including gene flow (Avise 2000; Knowles 2009; Hickerson et al 2010). New phylogeographic methods, and Bayesian phylogeography in particular, infer the geographic diffusion of a clade over time within a coalescent-based framework and have therefore enabled the simultaneous estimation of the spatial and temporal history of individuals and populations (Lemey et al 2009, 2010; Nylinder et al 2014). Whereas the initial implementation of Bayesian phylogeography required discretized areas (e.g., countries) and assumed a time-homogeneous process of geographic diffusion (Lemey et al 2009), recent modifications have enabled the analysis of continuous geographic data (e.g., latitude/longitude coordinates) and heterogeneous geographic diffusion rates amongst individuals, and most recently, amongst species (Nylinder et al 2014). However, examining species-level phylogeography requires an accurate knowledge of the species limits. But species limits, particularly within closely related groups of species in the tropics, are often unknown(e.g., Barley et al 2013). Identifying species in an objective manner is requisite to defining groups for species-level phylogeographic analysis.

The timing of sympatry or allopatry amongst ancestral ranges of closely related lineages can be determined by applying absolute dates to phylogeographic analyses. Knowing this information is of primary concern when comparing phylogeographic models of divergence with gene flow vs. a model of secondary contact. For instance, two species that presently have overlapping distributions might be assumed to be in secondary contact if the ancestral ranges of the species were allopatric (Pettengill & Moeller 2012). Similarly, determining colonization times in areas of hybridization can help define times of population expansion when testing models of gene flow (e.g., Smith et al 2011). And finally, understanding the geographic and temporal occurrence of particular clades along with the geologic history of the study region can help elucidate the biogeographic mechanisms shaping phylogeographic patterns (e.g., Chiari et al 2012).

Coalescent simulations are a valuable tool for testing alternative phylogeographic scenarios (e.g., Knowles 2001; Kuhner 2009; Pelletier & Carstens 2014). Modeling genetic variation within a coalescent framework enables quantitative tests of alternative population histories and the estimation of population genetic parameters (e.g., Hudson 2002). The modeled population histories are often generated based on inferences obtained from geological data (Carstens et al 2005), paleoclimatic data (Spellman & Klicka 2006), or based on previous genetic studies (Tsai & Carstens 2013), and the parameterizations used in the models can be derived from estimates from empirical data (Carstens et al 2005). The parameter estimates based on the empirical data are then compared to the distribution of simulated values, allowing for the support or rejection of each hypothesis. In such a way, a vast majority of phylogeographic models otherwise indistinguishable when only utilizing empirical data can be reduced to a reasonable set of candidate models (e.g., Pelletier & Carstens 2014).

In this study, we examined temporal and geographic patterns of gene flow to investigate the phylogeographic history of the *Sceloporus spinosus* group. The *S. spinosus* group consists of two species, *S. spinosus* and *S. horridus* (Wiens & Reeder 1997; Smith & Chiszar 1992) that are broadly distributed throughout xeric habitats in Mexico (Smith 1939; Cole 1970; Frost 1978). Each species is composed of three subspecies: *S. s. spinosus, S. s. apicalis,* and *S. s. caeruleopunctatus,* and *S. h. horridus, S. h. albiventris,* and *S. h. oligoporus* (Frost 1978; Smith & Chiszar 1992). *Sceloporus spinosus* is primarily found in and near the (eastern) Sierra Madre Oriental mountain range, whereas *S. horridus* is largely distributed in the lower slopes of the (western) Sierra Madre Occidental mountain range. Similarities in the habitat preferences of these two species have led to areas of sympatry, and suspected hybridization, in southern Mexico (Fig. 1).

**Figure 1:**
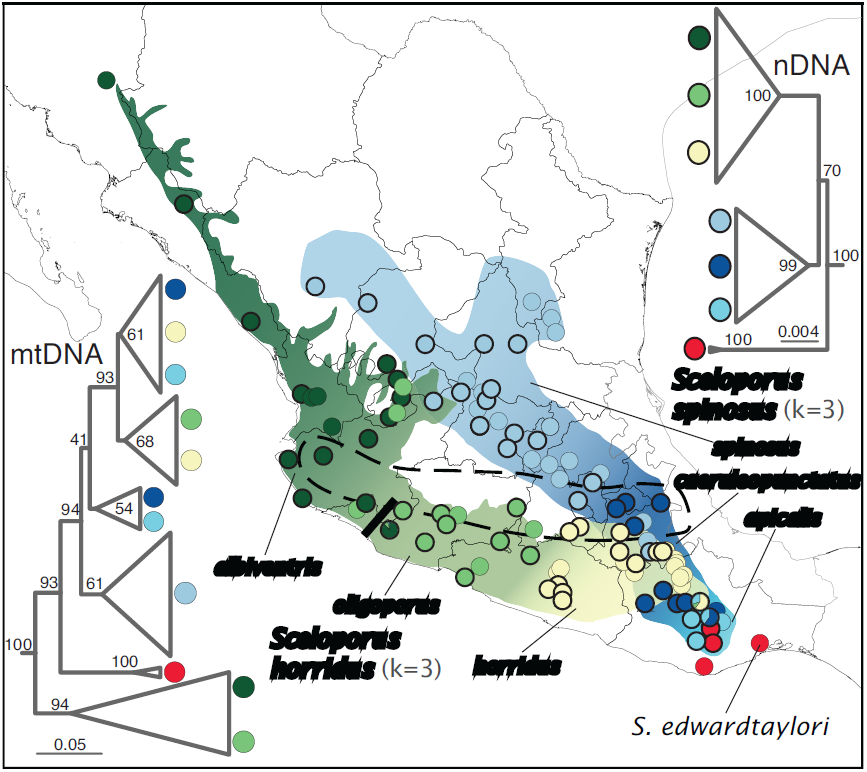
Sampling localities and species/subspecies distributions of *Sceloporus spinosus* and *S. horridus* in Mexico (based on (Smith 1939; Frost 1978; Smith & Chiszar 1992). Sampling localities with bold rings indicate specimens that have been amplified for nDNA in addition to mtDNA, whereas samples with a light ring have only mtDNA. The designation (i.e., color) of nDNA samples was based on Geneland assignments (3 inferred populations for each species), and the designation of mtDNA samples to subspecies was based on morphological characters. The Rio Ahuijullo Basin in southwestern Mexico is indicated with a dark line, and the approximate location of the Transvolcanic Belt is shown with a dashed line. Also shown are the concatenated mt- and nDNA trees inferred from RAxML, where values at nodes represent bootstrap (bs) proportions. Note the mixed clade of *S. horridus* and *S. spinosus* in the mtDNA tree, in addition to the contrasting phylogenetic placement of *S. edwardtaylori* between mt- and nDNA trees.

In addition to sharing habitat preferences, *S. spinosus* and *S. horridus* also share similar morphologies, thus making “attempts at determining phylogenetic relationships among *spinosus* group species on the basis of classical characteristics of scutellation and color pattern considerably frustrating” (Cole 1970). In fact, previous researchers have proposed contact zones in Puebla and Oaxaca to explain the observed morphological overlap in traits (Frost 1978; Boyer et al 1987). Beyond identification of species, distinguishing between subspecies has also proven to be difficult. For instance, overlap in quantitative characters exists between *S. spinosus* subspecies (Smith & Chiszar 1992), and intergradation has also been suspected between many of the subspecies (*S. s. spinosus* x *S. s. apicalis, S. s. apicalis* x *S. s. caeruleopunctatus,* and *S. h. albiventris* x *S. h. oligoporus*) (Frost 1978; Smith & Chiszar 1992).

We aim to determine the temporal and geographic extent of overlap between *S. spinosus* and *S. horridus* with multi-locus nuclear DNA (nDNA) and mtDNA. We first used population assignment and species delimitation analyses to identify the number and geographic boundaries of distinct populations within each species, and then inferred phylogenetic trees for the nDNA and mtDNA data. We then performed coalescent simulations to model potential historic phylogeographic scenarios that could have generated the strong pattern of mito-nuclear discordance that we observed in the empirical data. In addition to testing models of divergence with gene flow and secondary (2^°^) contact, we utilized a new Bayesian phylogeographic approach that estimates the diffusion of populations through time (Nylinder et al 2014). This approach provided us with temporal and spatial information for discriminating between models of divergence with gene flow vs. 2^°^ contact.

## Materials & Methods

### Taxon Sampling

One hundred fourty-eight individuals were sampled across the distributions of *S. horridus* and *S. spinosus* (Fig. 1; Supplemental Table 1). Four samples of *S. edwardtaylori* were included for analysis because recent work (using combined n- and mtDNA) showed that this taxon is nested within the *S. spinosus* group (Leaché 2010; Wiens et al 2010). Assignment of individuals to species and subspecies was based on morphological character descriptions by Smith (1939), Smith & Smith (1951), Frost (1978), and Smith & Chiszar (1992). Of these 152 individuals, 81 yielded nuclear sequence data. However, after data refinement (see below), a total of 70 individuals, including two *S. edwardtaylori* individuals, were represented in the nuclear DNA (nDNA) analyses. Both *S. horridus* and *S. spinosus* were nearly equally represented in the nDNA dataset (Supplemental Table 2). Three individuals of *Sceloporus clarkii* were included in our dataset to serve as the outgroup for phylogenetic analyses.

### Molecular Data Collection

Genomic DNA was extracted from tissue using the Qiagen extraction kit. A total of three mtDNA regions and eight nDNA loci were targeted for sequencing and analysis; five of the nDNA regions are protein-coding (BACH1, EXPH5, KIAA 2018, NKTR, and R35), one region is intronic (NOS1) and two are anonymous loci (for primer references, see (Rosenblum et al 2007; Townsend et al 2008; Portik et al 2012; Brandley et al 2011)). We sequenced portions of the mitochondrial genes encoding the fourth unit of the NADH dehydrogenase (ND4, and adjacent genes encoding the tRNAs for histidine, serine, and leucine; Arèvalo et al 1994), the 12S ribosomal gene (Leaché & Reeder 2002), and cytochrome B (Kocher et al 1989).

Standard PCR protocols were used to amplify mitochondrial DNA (mtDNA), whereas a “touch-down” protocol was used to amplify the nDNA regions (94^°^ C for 1:00, [0:30 at 94^°^ C, 0:30 at 61^°^ C, 1:30 at 68^°^ C] x 5 cycles, [0:30 at 94^°^ C, 0:30 at 59^°^ C, 1:30 at 68^°^ C] x 5 cycles, [0:30 at 94^°^ C, 0:30 at 57^°^ C, 1:30 at 68^°^ C] x 5 cycles, and [0:30 at 94^°^ C, 0:30 at 50^°^ C, 1:30 at 68^°^ C] for 25 cycles). Diploid nuclear genotypes were phased using the program PHASE (Stephens et al 2001) where alleles were discarded if any site probability was <0.95 (resulting in <20% data reduction). We tested for intragenic recombination using the difference in sum-of-squares test (McGuire & Wright 2000) in TOPALi (Milne et al 2009) using a step-size of 10bp and a window size of 100bp for 500 parametric bootstraps.

### Population Assignment

We explored two methods to identify distinct populations within the *S. spinosus* group that utilize multi-locus nDNA data and require no *a priori* knowledge of population assignment or number of populations. We used the Bayesian program STRUCTURAMA (Huelsenbeck & Andolfatto 2007; Huelsenbeck et al 2011) to identify the number of populations (k) present in our data. Under a Dirichlet process prior, this program assigns individuals to populations while allowing both the allocation of individuals to populations and number of populations to be random variables. This assumes that the joint prior probability on the number of populations and allocation of individuals to populations follows a Dirichlet process prior, and the prior on the number of populations (k) can be specified. We chose the mean value for the prior distribution on k to range between 1 and 10 to ensure that the number of populations inferred was not sensitive to the prior mean value for the number of populations (k), assuming no admixture between populations (assuming admixture resulted in unstable results, where more populations were inferred than individuals in our dataset). The input data for STRUCTURAMA analyses were the two alleles with the highest posterior probabilities at each locus from our PHASE analyses. We ran four replicates of each STRUCTURAMA analysis for a length of 2x10^6^ generations and a burn in of 4x10^5^. We present results as the arithmetic mean of the four replicate analyses.

To estimate the number of populations within a geographic context, we used the program Geneland (Guillot et al 2005a,b; Guillot 2008). This program uses a spatial statistical model and Markov chain Monte Carlo sampling with GPS coordinates and multi-locus genotypes to estimate the number of populations, individual assignment probabilities, and the geographic limits between populations that are in Hardy-Weinberg equilibrium. Geneland utilizes the colored Poisson-Voronoi tessellation model to determine the unknown number of polygons that approximate the pattern of population spread over space, where the number of polygons follows a Poisson distribution. We varied the number of populations from 1-10 with a spatial correlation between allele frequencies, and ran five independent analyses with the same parameters for 10^6^ generations and a burn-in of 2x10^5^ generations. The spatial correlation model is more powerful at detecting subtle population differentiation (over the un-correlated model), and corresponds to the spatial patterns that can be expected when differentiation occurs by limited gene flow produced by physical barriers such as roads, rivers, or mountain ranges. We modified the format of the Geneland output files and combined the results with the program CLUMPP (Jakobsson & Rosenberg 2007) to generate individual assignment probabilities.

### Phylogenetic Tree Estimation

We estimated maximum likelihood phylogenetic trees for each nDNA locus in addition to concatenated nDNA and mtDNA datasets separately to examine the concordance in evolutionary history between these genomes. RAxML (Stamatakis 2006) was used with the GTR + Γ nucleotide substitution model and run for 500 nonparametric bootstrap iterations for both n- and mtDNA analyses, where one out of two alleles was randomly chosen to represent each individual in the concatenated nDNA analysis. Partitioning the data by gene vs. codon position did not affect topology or branch length estimates, so we present results from partitioning by codon position. We ran two replicates of each analysis to ensure stability of our results. Phylogenetic relationships were considered significant when bootstrap (bs) values were > 70% (Hillis & Bull 1993; Alfaro et al 2003).

### Species Delimitation

To delimit evolutionarily independent lineages, we performed Bayes Factor Delimitation of species (BFD) using only the nDNA dataset (Grummer et al 2014). Our species delimitation models were based on a combination of the results from population assignment and migration analyses. Gene flow violates the coalescent model used in the species tree estimation program *BEAST (Heled & Drummond 2010), so we performed species delimitation on two distinct datasets in an effort to remove potentially admixed individuals located on population margins. One dataset consisted of individuals limited to the “core” range of each population as determined in Geneland (see Results section below), whereas the other dataset consisted of all individuals (Fig. 3). Our expectation was that the dataset consisting of all individuals would be more likely to support the recognition of fewer species because under certain migration conditions (e.g., recent migration at a high rate), it is possible for gene flow between populations to homogenize gene pools and make divergent populations appear as one.

Six species delimitation models were tested against each other with each dataset: 1) the six-population model where each population based on population assignment analyses was distinct (the “6 pop” model), 2) a model of five species where the northern and central populations of *S. horridus* were lumped together(the “northern horridus migration” model), 3) a second five-species model where central and southern populations of *S. horridus* were lumped together (the “southern horridus migration” model), 4) a third five-species model with central and southern populations of *S. spinosus* lumped together (the “southern *spinosus* migration” model), 5) a four-species model with all populations of *S. horridus* lumped together (the “all *horridus* migration” model), and lastly 6) a two-species model where the three populations of each *S. horridus* and *S. spinosus* are represented as a single species (the “2 pop” model). Models 2-5 are based on “lumping” lineages together that were inferred to have non-zero migration rates between them (see Results below). We estimated species trees for these species delimitation models in *BEAST v1.7.5 (Heled & Drummond 2010) with the 8-locus nuclear dataset, where individuals were assigned to lineages based on our population assignment and BFD results (see Results section below). Species tree analyses only included individuals that did not show signs of admixture (i.e., we only included individuals with >0.90 posterior probability for belonging to one population). Each gene was given its own partition and analyzed under the uncorrelated lognormal molecular clock with the preferred substitution model as mentioned above. Analyses were run for 3x10^8^ generations, logging every 2x10^4^ steps, and convergence was assessed in Tracer v1.5 (Rambaut & Drummond 2007).

We selected the best species delimitation model through Bayes factor (Bf) analysis of the path sampling (“PS”) and stepping stone (“SS”) marginal likelihood estimates (Baele et al 2012; Grummer et al 2014). Previous research has shown that PS and SS marginal likelihood estimates are much more accurate at estimating the marginal likelihoods of models over harmonic mean estimation (Fan et al 2011; Xie et al 2011; Baele et al 2012; Grummer et al 2014). Briefly, PS and SS marginal likelihood samplers estimate the marginal likelihood in a series (either a continuous path or a broken path of “stepping stones”) that bridges the posterior and prior distribution of a model. In this way, the influence of the prior information is accounted for and the marginal likelihood is not overestimated. All models were compared against each other within the two datasets (“All Samples” and “Core Samples”), and the top model is considered significantly better than the rest if the Bf value (= twice the difference in marginal likelihood estimates) is greater than 10 (Kass & Raftery 1995).

### Temporal Estimation of Gene Flow

We performed simulations to discern whether gene flow occurred amongst ancestral (i.e., divergence with gene flow) or extant populations (i.e., secondary contact). To determine when gene flow occurred in the *S. spinosus* group, we used the genealogical sorting index (gsi; Cummings et al 2008). The gsi is a statistic that estimates the degree of exclusive ancestry of individuals in labeled groups on a rooted tree and is a statistically more powerful measure of population divergence than F_*ST*_ (Cummings et al 2008). The gsi statistic can range from 0 to 1, where the maximum value of 1 is achieved when a group is monophyletic, and is normalized to account for disparities in group sizes while also accommodating unresolved relationships (i.e., polytomies). Although genealogical exclusivity is a function of the sorting of ancestral polymorphisms, allele sharing could also be due to the extent and timing of migration events. We therefore modeled migration scenarios and performed coalescent simulations to test models of divergence with gene flow vs. 2^°^ contact, which have explicit expectations about the timing of migration events.

Coalescent simulations were performed in the program MCcoal (Rannala & Yang 2003; Yang & Rannala 2010). In our simulations, we used a symmetric migration matrix and held the migration rate constant at 1 *N*_*e*_*m* (0.5 *N*_*e*_*m* in each direction, where *N*_*e*_*m* equals the product of the effective population size and the migration rate per generation), but varied the migration start and end times (Fig. 2). We used a migration rate of 1 *N*_*e*_*m* because this is the maximum rate of migration allowed between populations until they are considered separate species by some researchers (Porter 1990; Hey 2009). We therefore consider this rate a minimum when modeling interspecific migration. Divergence times and population sizes used in the simulations were derived from estimates of our empirical data in the programs BP&P (Yang & Rannala 2010) and Arlequin v3.5 (Excoffier & Lischer 2010), respectively. We simulated species trees including no gene flow (Scenario A; Fig. 2a), ancestral gene flow between the common ancestors of *S. horridus* and *S. spinosus* (Scenario B; Fig. 2b), gene flow between ancestral populations as well as contemporary gene flow between one *S. horridus* and two *S. spinosus* lineages (lineages selected based on empirical results, see Results; Fig. 2c), gene flow between the common ancestors of *S. horridus* and *S. spinosus*, followed by a cessation of gene flow until contemporary gene flow between three lineages as above (Scenario D; Fig. 2d), and contemporary gene flow between one lineage of *S. horridus* and two lineages of *S. spinosus* (Scenario E; Fig. 2e). We restricted our simulations of gene flow to these models because the mtDNA clade showing admixture was comprised only of individuals from these three populations.

**Figure 2:**
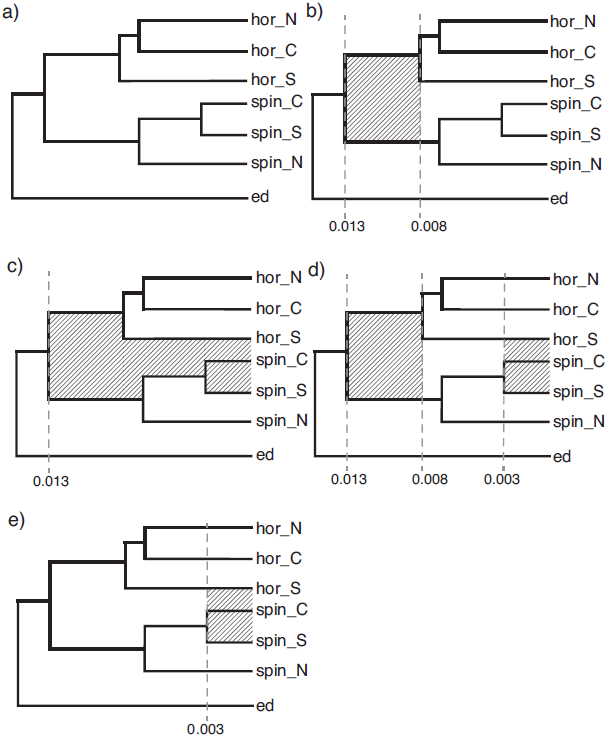
The six scenarios modeled for coalescent-based simulations. Migration times are indicated by the species divergence time parameter *τ* (= expected number of mutations per site), and migration events are indicated by diagonal shading. Northern, central, and southern populations are denoted by “N”, “C”, and “S”, respectively.

We simulated 10,000 gene trees under each model, then calculated a gsi value for each group within each gene tree in the “genealogicalSorting” R package (using the “multitree” function). We focused empirical gsi calculations on the mtDNA because this locus showed signs of admixture at clade boundaries. Furthermore, lower resolution in the nDNA led to high variability in GSI values for each nuclear locus that were difficult to interpret. To account for phylogenetic uncertainty in the empirical data, we calculated a single ensemble gsi value (a weighted sum of gsi values from each tree in the posterior distribution) for the mtDNA for each population on a posterior distribution of 8,000 trees inferred in MrBayes (v3.2; Ronquist and Huelsenbeck 2003). For the MrBayes analysis, we partitioned the dataset by codon for protein-coding genes (one 12S partition, three partitions each for CytB, and four partitions for ND4 including the tRNA coding sequence) and assigned each the best substitution model determined in jModelTest v2 (Darriba et al 2012; Guindon & Gascuel 2003). We ran two analyses for 10^7^ generations, sampling every 2000 steps, and discarded the first 20% as burn-in (determined by visual examination in Tracer v1.5 Rambaut & Drummond 2007).

To assess the probability that the empirical mtDNA gsi values are different from the gsi values from the simulated trees, we calculated the frequency of simulated gsi values that were in the tail of the distribution beyond the empirical value; we compared all gsi values from the simulations to our empirical dataset (e.g., all simulation values were “accepted”). These values could therefore be interpreted as one-half of the p-value statistic when testing the null expectation that the empirical mtDNA gsi values were drawn from the simulated gsi distribution. The comparison of empirical mtDNA gsi values to the simulated gsi values provide a statistical test of determining the timing of migration events in the *S. spinosus* group.

### Estimation of Nuclear Gene Flow

We estimated ancestral and contemporary levels of gene flow in the program IMa2 (Hey 2010) using our empirical nDNA. This program estimates bi-directional and uni-directional migration rates, divergence times, and population sizes. The IM model assumes non-recombinant loci, constant population sizes, and that population-level sampling has been performed randomly. We did not include the mtDNA in this analysis because of the difficulty in assigning mtDNA haplotypes to nuclear-based species that are paraphyletic in the mtDNA tree. We performed analyses on three separate datasets, where the user-specified topologies were based on our empirical species tree estimate (see below): (1) only *S. horridus* populations (=3 extant populations), (2) only *S. spinosus* populations (=3 extant populations), and (3) both *S. horridus* and *S. spinosus* (=6 extant populations). For the three-population models, we specified 3x10^5^ steps as burn-in with 3x10^5^ steps following burn-in, and allowed the program to infer migration rates amongst all pairwise lineage combinations (including ancestral gene flow). For the 6-population model, the burn-in period lasted for 5x10^5^ generations followed by 3x10^5^ steps post burn-in, and we estimated migration between all pairwise lineage combinations (including ancestral gene flow). Whereas the three-population models allowed us to examine gene flow between populations within each species (including ancestral gene flow), the 6-population model enabled us to test for gene flow *across* species (both extant and ancestral lineages). For all models, we ran four replicate analyses (using different starting seeds) of 100 chains with heating terms of 0.98 and 0.90 (options -ha and -hb), and a maximum value of five on the uniform prior for the migration value. Convergence of independent runs was confirmed by examining trace plots for stationarity and ESS values of all estimated parameters (all ESS values were > 5000). As estimated migration values across runs were highly similar, we report the results here from a single analysis. Significant levels of migration were assessed using the Nielsen & Wakeley (2001) test implemented in IMa2.

### Bayesian Phylogeographic Analysis

We utilized Bayesian phylogeography (Lemey et al 2009, 2010) to determine the temporal and geographic extent of overlap, and therefore the possibility of introgression, between the populations comprising the “admixed” mtDNA clade. We utilized a method that was recently developed by Nylinder et al (2014) that applies the relaxed random walk (RRW) continuous phylogeographic approach (Lemey et al 2010) to relax the assumption of geographic rate diffusion homogeneity across branches in the species tree. This method follows a two-tiered approach in the program BEAST (Drummond & Rambaut 2007) where a posterior distribution of species trees is first generated, which is then subsequently used in an RRW analysis. To generate the species tree, we used *BEAST v1.7.5 (Heled & Drummond 2010) on the 8-locus nuclear dataset as described in the BFD section above. We calibrated the root of the (*S. spinosus* group + *S. edwardtaylori*) clade at 5.0 million years ago (mya) with a standard deviation of 0.5, based on the time-calibrated tree from Leaché & Sites Jr (2010).

The species tree diffusion analysis was performed with BEAST v1.8.1. We used LogCombiner v1.8 from the BEAST package to combine and thin results from three independent species tree analyses. We pruned *S. edwardtaylori* in the program Mesquite (v2.75; Maddison and Maddison 2011) because the phylogeographic history of this species was not the focus of this study, and then used one thousand species trees from the posterior distribution as input for the species tree diffusion analysis. We circumscribed polygons in Google Earth to approximate extant distributions for each lineage/population based on published range maps (Smith 1939; Frost 1978; Smith & Chiszar 1992) and Geneland results; these polygons were then referenced along with the posterior distribution of species trees for analysis. The RRW process rescales the precision matrix of the diffusion process by a branch-specific scalar drawn from, in this case, a lognormal distribution centered on 1.0. As in Nylinder et al (2014), a prior exponential distribution on the standard deviation of the lognormal distribution was specified with a mean of 2.712. We explored the effect of (geographic) starting location on species-level geographic diffusion by choosing two different starting locations within each species’ boundaries. All priors on the RRW diffusion model were kept the same as in Nylinder et al (2014), to which we refer the reader for further details on this method. We ran four independent replicates of species tree diffusion analysis for 5x10^8^ generations each, logging every 5x10^5^ generations. The “time slice” function of the program SPREAD (Bielejec et al 2011) was then used to visualize the ancestral 80% HPD regions in Google Earth at 5x10^5^ year intervals from 3.0 - 0.5mya. All files used for Bayesian phylogeographic analysis are available online as supplementary materials and in the Dryad digital online repository.

## Results

### Taxon Sampling

We generated mtDNA data for 74 *S. horridus*, 74 *S. spinosus*, and four *S. edwardtaylori* (Supplemental Table S1). Our nDNA dataset consisted of a subset of the individuals present in the mtDNA dataset: 36 *S. horridus*, 32 *S. spinosus*, and two *S. edwardtaylori* (Supplemental Table S2). All individuals in the mtDNA dataset were amplified for at least one of the three mitochondrial regions examined, whereas the final nDNA dataset only consisted of individuals with sequence data for ≥ 4 loci (≥ 50% complete matrix) due to poor genomic DNA quality for particular individuals at some loci.

### Molecular Data Collection

The three mtDNA regions varied in length from 782-1025bp and totaled 2,639bp with 859 variable sites, 714 of which were parsimony-informative (Table 1; GenBank accession nos. xxxx-xxxx). In contrast, the eight nDNA regions ranged from 485-1247bp and totaled 5,716bp with 459 variable sites and 420 parsimony-informative sites (GenBank accession nos. xxxx-xxxx). Large indels (>10bp) were present in the intron (NOS1) and two anonymous loci (Sun_035, Sun_037), but these were not scored for usage in the phylogenetic analyses. No evidence of intra-genic recombination was detected in any gene.

**Table 1:** Information for the genetic data gathered in this study. The first three regions are mitochondrial regions, whereas the remainder are nuclear regions. Gene region “Type” abbreviations indicate noncoding (NC), protein-coding (PC), intron (I), and anonymous (A).

| Gene<br>Region | Type | Length<br>(bp) | Variable<br>Sites | Parsimony-Informative<br>Sites | DNA<br>Substitution<br>Model |
| --- | --- | --- | --- | --- | --- |
| 12S | NC | 782 | 141 | 113 | HKY+I+$\Gamma$ |
| CytB | PC | 1025 | 412 | 352 | HKY+I |
| ND4 | PC | 832 | 306 | 249 | HKY+I+$\Gamma$ |
| BACH1 | PC | 1247 | 91 | 83 | HKY+I |
| EXPH5 | PC | 900 | 55 | 49 | HKY+$\Gamma$ |
| KIAA | PC | 621 | 27 | 25 | HKY+I |
| NKTR | PC | 617 | 54 | 48 | HKY+I |
| NOS1 | I | 666 | 68 | 66 | HKY+$\Gamma$ |
| R35 | PC | 658 | 43 | 35 | HKY+I |
| Sun_035 | A | 522 | 49 | 46 | HKY+I |
| Sun_037 | A | 485 | 72 | 68 | HKY+$\Gamma$ |
| Total | — | 8,355 | 1,318 | 1,134 | — |

### Population Assignment

Under the “no admixture” model, STRUCTURAMA identified six populations based on the nDNA dataset (Table 2). When the prior mean on k was ≥7, seven populations were inferred, indicating some sensitivity of our analysis to the prior distribution on k. Geneland results provided support for six distinct populations, where this model (k=6) received >0.65 of the posterior probability. Three of these populations were composed of *S. horridus* individuals, and the other three populations were composed of *S. spinosus* individuals (Fig. 3). Proportions of population assignment based on Geneland output are shown in Figure 1. Nearly all individuals (65/68) showed >0.95 probability in belonging to a single cluster. The geographic boundaries of the populations inferred in Geneland are largely in agreement with currently recognized subspecific boundaries.

**Table 2:** Results from STRUCTURAMA indicating the posterior probability values when the prior mean on the number of populations (k) was varied. Values shown are the average of four independent runs. Bold values indicate the highest posterior probability for each mean value on the prior for k. See Materials & Methods section for further details on this method.

| Number of<br>Populations<br>Inferred | Prior Mean on Number of Populations (k) |  |  |  |  |  |  |  |  |  |
| --- | --- | --- | --- | --- | --- | --- | --- | --- | --- | --- |
|  | 1<sup>1</sup> | 2 | 3 | 4 | 5 | 6 | 7 | 8 | 9 | 10 |
| 1 | 0.00 | 0.00 | 0.00 | 0.00 | 0.00 | 0.00 | 0.00 | 0.00 | 0.00 | 0.00 |
| 2 | 0.00 | 0.00 | 0.00 | 0.00 | 0.00 | 0.00 | 0.00 | 0.00 | 0.00 | 0.00 |
| 3 | 0.00 | 0.00 | 0.00 | 0.00 | 0.00 | 0.00 | 0.00 | 0.00 | 0.00 | 0.00 |
| 4 | 0.00 | 0.01 | 0.00 | 0.00 | 0.00 | 0.00 | 0.00 | 0.00 | 0.00 | 0.00 |
| 5 | 0.00 | 0.21 | 0.10 | 0.05 | 0.04 | 0.02 | 0.01 | 0.01 | 0.01 | 0.01 |
| 6 | 0.00 | **0.68** | **0.67** | **0.60** | **0.52** | **0.45** | 0.38 | 0.31 | 0.27 | 0.22 |
| 7 | 0.00 | 0.10 | 0.21 | 0.30 | 0.37 | 0.41 | **0.45** | **0.46** | **0.46** | **0.45** |
| 8 | 0.00 | 0.00 | 0.02 | 0.05 | 0.07 | 0.11 | 0.15 | 0.19 | 0.22 | 0.26 |
| 9 | 0.00 | 0.00 | 0.00 | 0.00 | 0.01 | 0.01 | 0.02 | 0.03 | 0.05 | 0.06 |
| 10 | 0.00 | 0.00 | 0.00 | 0.00 | 0.00 | 0.00 | 0.00 | 0.00 | 0.01 | 0.01 |

<sup>1</sup> Results from the analysis with a k prior of 1 were unstable and reported a posterior probability of 1.0 for 68 populations

**Figure 3:**
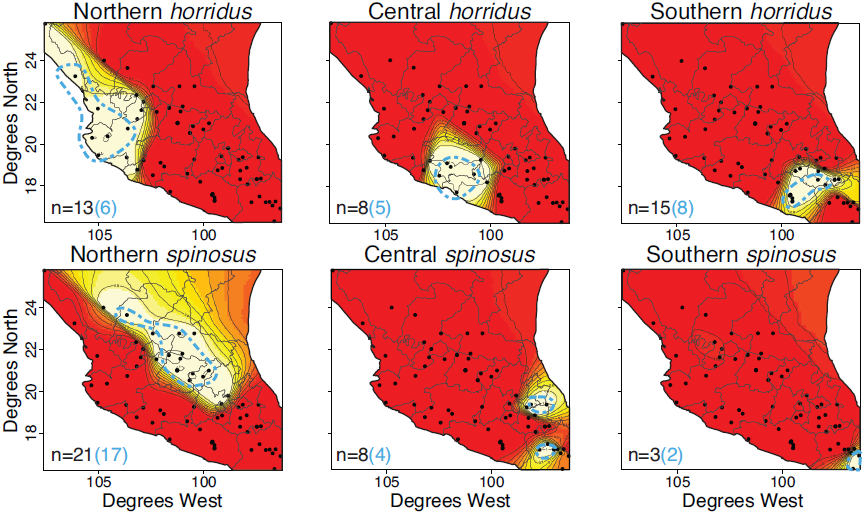
Geneland analysis results showing the number of populations and the probability of individual assignment to each population. These results can be interpreted as topographic maps, where white colors indicate high probabilities of assignment to that cluster and red represents low assignment probability. Blue dashed lines indicate which samples were included in the “core” sampling for BFD analyses. Black numbers in the lower left portion of each tile are the number of individuals in that cluster, whereas the blue number represents the number of individuals from that cluster in the “core” sampling scheme.

### Phylogenetic Tree Estimation

Phylogenetic trees for six out of the eight nDNA loci revealed moderate to strong support for the monophyly of one species to the exclusion of the other, whereas the remaining two loci showed some degree of species-level paraphyly (Supplemental Fig. 1). Support values towards the tips of the trees (i.e., between alleles) were generally low. The position of *S. edwardtaylori* was variable across gene trees. The concatenated nDNA tree revealed strong support (bs = 100) for the sister relationship between *S. edwardtaylori* and (*S. spinosus + S. horridus*) (Figs. 1,4; Supplemental Fig. 2). The support for mutual exclusivity between *S. spinosus* and *S. horridus* was strong with bs values of 99 and 100 for each group, respectively. The nDNA was geographically structured with strongly supported clades in general agreement with currently recognized subspecific geographic boundaries (Fig. 4). However, it is important to note that not all populations inferred in our population assignment tests appear as natural groups in the nDNA concatenated tree, specifically, central *S. horridus* and southern *S. spinosus* (Fig. 4).

**Figure 4:**
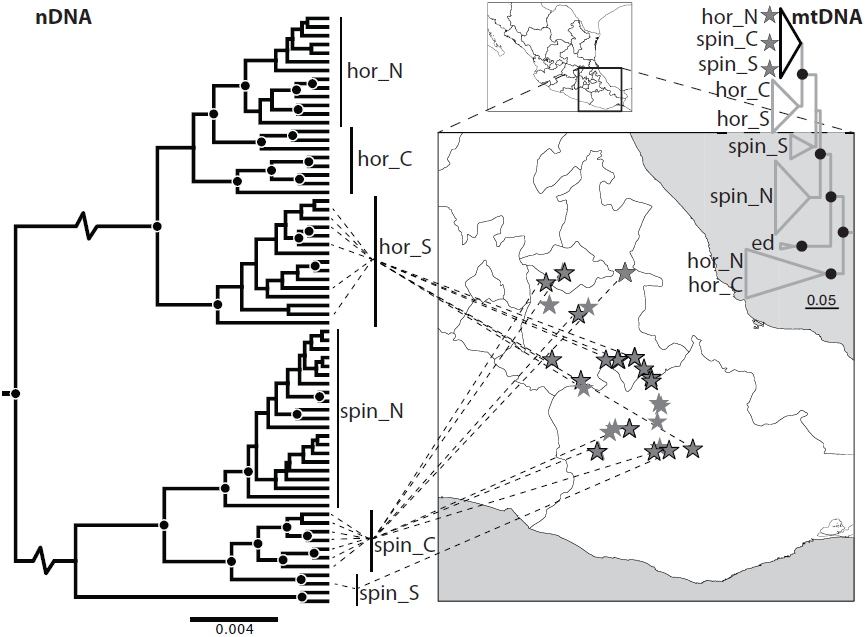
Map showing geographic locations of putatively admixed individuals in southeastern Mexico along with their phylogenetic positions in n- and mtDNA trees. Stars without bold outlines indicate individuals without nDNA data, and black dots in the phylogenetic trees indicate bootstrap values >70. Note that groupings identified on the nDNA tree are based on population assignment analyses and therefore are not all monophyletic groups (e.g., central *S. horridus* and southern *S. spinosus*).

The (*S. edwardtaylori*, (*S. spinosus, S. horridus*)) relationship inferred with the nDNA is in stark contrast to the relationships inferred with the mtDNA. In the mtDNA tree, both *S. spinosus* and *S. horridus* were paraphyletic, with S. edwardtaylori nested within these two species with strong support (bs = 93; Figs. 1,4; Supplemental Fig. 3). The mtDNA tree also shows two clades of *S. spinosus* and two clades of *S. horridus*, in addition to one moderately supported clade (bs = 61) consisting of both *S. spinosus* and *S. horridus* individuals. Interestingly, the *S. spinosus* and *S. horridus* individuals in this “admixed” clade occur in southern Mexico where these two species are sympatric (Fig. 4). Although these putatively admixed individuals form a clade in the mtDNA tree, they belong to three distinct populations in the nDNA, specifically, central *S. spinosus*, southern *S. spinosus*, and southern *S. horridus* (Fig. 4). This phylogeographic result was interpreted as geographically localized mitochondrial introgression, as incomplete lineage sorting in the mtDNA would be expected to not leave a strong geographic signature. We therefore performed coalescent simulations with gene flow and used the gsi statistic to determine the timing of this admixture.

### Species Delimitation

Marginal likelihood estimates based on both PS and SS marginal likelihood estimators were very similar, and the ranking of models was identical, so we therefore only show the PS results. Out of the six species delimitation models examined, the model containing six species (corresponding to the six populations identified through population assignment analyses) was favored over all other models by a Bf > 70 (Table 3). This result was consistent across both datasets composed of all samples and “core” samples. These results did not match our expectation, given that non-zero levels of gene flow were detected between three population-pairs (see “Estimation of Nuclear Gene Flow” results below). The “2 pop” model that represented *S. horridus* and *S. spinosus* each as a single species composed of three populations was the lowest ranked model in both datasets, indicating the strong possibility that currently described subspecies may warrant the recognition as distinct species.

**Table 3:** Results from Bayes Factor Delimitation of species (BFD) analyses. Path sampling (“PS”) and stepping stone marginal likelihood estimates were very similar, so we only show the PS results here. The Bayes Factor value represents two times the difference in marginal likelihood estimates between each model and the top model (“6 pop”). See Materials and Methods section for the composition of each species delimitation model.

| Model | # Species | All Samples |  | “Core” Samples |  |
| --- | --- | --- | --- | --- | --- |
|  |  | PS | Bayes Factor | PS | Bayes Factor |
| 6 pop | 6 | -12854 | — | -11353 | — |
| southern *spinosus* migration | 5 | -12889 | 71 | -11396 | 86 |
| northern *horridus* migration | 5 | -12971 | 234 | -11425 | 144 |
| southern *horridus* migration | 5 | -12978 | 248 | -11428 | 151 |
| all *horridus* migration | 4 | -13181 | 654 | -11536 | 367 |
| 2 pop | 2 | -13419 | 1129 | -11714 | 721 |

### Temporal Estimation of Gene Flow

We focus our gsi results on the central *S. spinosus*, southern *S. spinosus*, and southern *S. horridus* populations (and their ancestors), because these populations appeared to be admixed in the mtDNA tree and therefore were the populations in which we modeled gene flow (see Figs. 2,4). When gene flow was not modeled in our simulations, gsi values were relatively high (all values ≥ 0.66; Scenario A; Table 3), i.e., relatively high levels of monophyly within populations. The gsi values reported for Scenario B, which included only historic gene flow between the common ancestors of *S. horridus* and *S. spinosus* (and therefore represents the model of divergence with gene flow), were similar to (but all less than) those reported for the model with no gene flow (Scenario A; Table 3; Supplemental Fig. 4), indicating that the gsi index did not do well at detecting ancestral gene flow. When migration amongst extant populations was included in the model (e.g., Scenarios C-E), gsi values markedly decreased (Table 3; Supplemental Fig. 4), particularly for the populations in which migration was modeled, demonstrating that the gsi statistic does much better at detecting recent gene flow, as opposed to ancestral gene flow.

Based on the empirical mtDNA data, central and southern populations of *S. horridus* along with the central *S. spinosus* population returned the lowest gsi values (< 0.55; Table 3), whereas gsi values for the other populations were all ≥ 0.90 (Table 3). According to our test statistic, the probability that southern *S. horridus* had a history similar to those modeled by Scenarios A and B is very low (0.0002, and 0.003, respectively), meaning that this population experienced appreciable levels of ancestral gene flow (> 1*N*_*e*_*m*; Fig.2; Table 3). However, there is strong probability that central and southern *S. spinosus* populations match the history of Scenarios A and B (all p≥0.09 for rejecting these scenarios), meaning they experienced negligible levels of ancestral gene flow (<1*N*_*e*_*m*; Table 3). The empirical mtDNA gsi values for southern *S. horridus* and central *S. spinosus* populations strongly matched the simulated distribution values (all p>0.11 for rejecting these scenarios) when gene flow was modeled amongst extant lineages (Scenarios C-E; Figs. 2,5; Table 3). However, the empirical gsi value for the southern *S. spinosus* population did not fit the expected distribution of simulated gsi values (p<0.03 for rejecting these scenarios) resulting from these same scenarios modeling recent gene flow (Fig. 5; Table 3).

**Figure 5:**
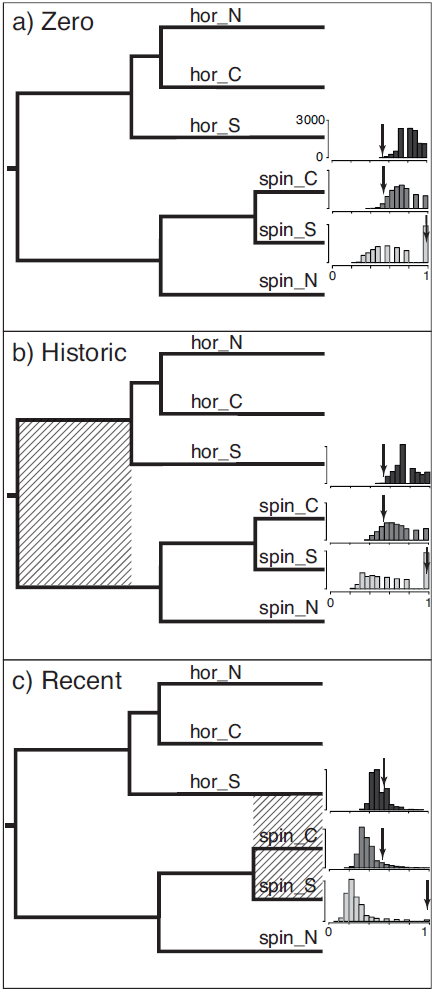
Gsi (genealogical sorting index) results for both simulated and empirical datasets along with the species tree topology used in the simulations. Histograms to the right indicate the distribution of gsi values recorded during simulations (see text for simulation details) for central *Sceloporus spinosus*, southern *S. spinosus*, and southern *S. horridus*. Y-axis values range from 0-3000, and x-axis values of the gsi statistic range from 0-1. Black arrows indicate the gsi value for the mtDNA empirical data. Figure (a) shows the gsi results for the model with no migration (Scenario A), (b) represents historic gene flow only (Scenario B), and (c) represents the gsi values for Scenario E that models recent gene flow (histograms for Scenarios C,D looked nearly identical).

### Estimation of Nuclear Gene Flow

Although the 3-population models are nested subsets of the 6-population model, the IMa2 results were inconsistent between these analyses (Table 4). Significant levels of unidirectional gene flow were detected within *S. horridus*, from northern *S. horridus* into southern *S. horridus*, from southern *S. horridus* into central *S. horridus*, and historically, between the common ancestor of northern and central populations *S. horridus* populations with the southern *S. horridus* population (Table 4; Supplemental Fig. 5). Within *S. spinosus*, significant levels of gene flow were detected from the central *S. spinosus* population into southern *S. spinosus*, and historically, from northern *S. spinosus* into the common ancestor of central and southern *S. spinosus* populations (Table 4). The full 6-population model allowed us to test for gene flow between *S. horridus* and *S. spinosus*. In terms of migration across species, a significant migration rate was reported from southern *S. horridus* into northern *S. spinosus*, a result coincident with a scenario of interspecific mitochondrial introgression (Table 4). We tested whether or not these patterns were the result of isolation-by-distance through a Mantel test in the program Alleles in Space (for each species separate and combined; (Miller 2005)), and the results were insignificant (results not shown).

**Table 4:** Gsi values for both empirical (mtDNA) and simulated datasets for all scenarios modeled (see Fig. 2). Northern, central, and southern populations are denoted by “N”, “C”, and “S”, respectively. Numbers in parentheses indicate the frequency of simulation results more extreme than the empirical gsi value.

| Lineage | Empirical |  | Simulations |  |  |  |
| --- | --- | --- | --- | --- | --- | --- |
|  | mtDNA | Scenario A | Scenario B | Scenario C | Scenario D | Scenario E |
| N *horridus* | 0.90 | 0.81 | 0.76 | 0.76 | 0.77 | 0.81 |
| C *horridus* | 0.37 | 0.78 | 0.74 | 0.74 | 0.74 | 0.78 |
| S *horridus* | 0.54 | 0.82 (0.0002) | 0.76 (0.003) | 0.50 (0.248) | 0.50 (0.262) | 0.50 (0.245) |
| N *spinosus* | 1.00 | 0.93 | 0.91 | 0.90 | 0.91 | 0.93 |
| C *spinosus* | 0.53 | 0.73 (0.045) | 0.67 (0.216) | 0.39 (0.059) | 0.39 (0.063) | 0.39 (0.072) |
| S *spinosus* | 0.97 | 0.66 (0.289) | 0.63 (0.286) | 0.27 (0.010) | 0.27 (0.011) | 0.27 (0.011) |

**Table 5:**
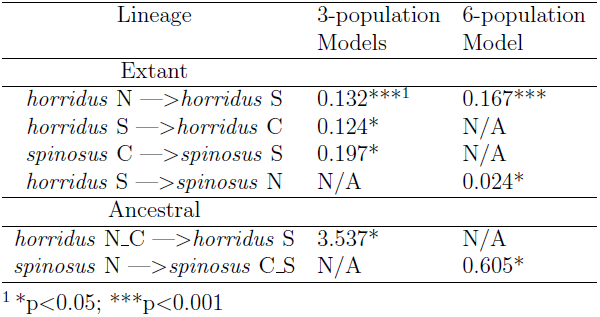
Significant results from the isolation-migration (IMa2) analyses. Values given are in *2Nm* per generation, and N/As indicate that no significant migration estimates were reported for that model (e.g., 3-population or 6-population model). Northern, central, and southern populations are denoted by “N”, “C”, and “S”, respectively. Common ancestors of two lineages are indicated with an underscore (_) between daughter lineage population numbers (e.g., *horridus* N_C is the common ancestor of northern and central *S. horridus* populations). Asterisks indicate significance levels for the Nielsen & Wakeley (2001) test.

| Lineage | 3-population<br>Models | 6-population<br>Model |
| --- | --- | --- |
| Extant |  |  |
| *horridus* N —> *horridus* S | 0.132***<sup>1</sup> | 0.167*** |
| *horridus* S —> *horridus* C | 0.124* | N/A |
| *spinosus* C —> *spinosus* S | 0.197* | N/A |
| *horridus* S —> *spinosus* N | N/A | 0.024* |
| Ancestral |  |  |
| *horridus* N_C —> *horridus* S | 3.537* | N/A |
| *spinosus* N —> *spinosus* C_S | N/A | 0.605* |

<sup>1</sup> \*p<0.05; \*\*\*p<0.001

### Bayesian Phylogeographic Analysis

Only one (*S. h. horridus*) individual was removed for the species tree analysis due to an admixed genotype (Fig. 1). The time-calibrated species tree revealed a root age of 3.1 mya (1.55-5.61 95% C.I.) for the *S. spinosus* group (results not shown). Altering the starting coordinates for each population did not appear to have an affect on our species tree diffusion analyses. At 3.0 and 2.5 mya, the distributions of the common ancestors (CA) of *S. horridus* and *S. spinosus* were largely sympatric in southern Mexico (Fig. 6). *Sceloporus horridus* split into two lineages at 2.1 mya, where southern *S. horridus* was nearly 100% sympatric with the *S. spinosus* CA. At 1.5 mya, southern *S. horridus* had moved slightly to the east and shares less range overlap with the CA of *S. spinosus*. By 1.0 mya, southern *S. horridus* and the CA of central and southern *S. spinosus* populations overlap with each other by approximately 60%. At 0.5 mya, the central *S. spinosus* population is nearly 100% sympatric with the southern *S. spinosus* population, and southern *S. horridus* is sympatric in the east with both central and southern populations of *S. spinosus* (Fig. 6). These results indicate that all populations present in the admixed mtDNA clade were largely sympatric throughout their existence until within the past one million years, at which point populations began diverging in allopatry.

**Figure 6:**
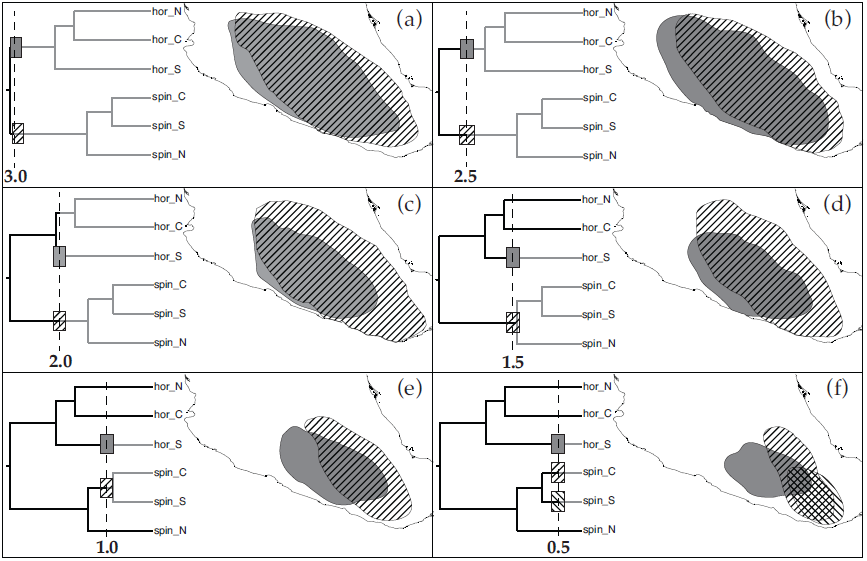
Bayesian phylogeographic results under the relaxed random walk (RRW) species tree diffusion approach. Distributions indicate the 80% HPD location of the depicted lineages from 3.0 (a) to 0.5 mya (f) using the “time slice” feature in SPREAD.

### Discussion

Recent analytical advancements in gene flow detection have given researchers the ability to utilize multi-locus datasets to estimate migration not only amongst extant lineages, but also between ancestral lineages (e.g., Hey 2010). Similarly, phylogeographic analyses can be tested in a statistical framework (e.g., Chan et al 2011; Pelletier & Carstens 2014). However, identifying the extent of historic geographic overlap and/or separation of lineages, parameters critical to differentiating between secondary contact and divergence with gene flow, has remained difficult (e.g., Pettengill & Moeller 2012). In this study, we employed phylogeographic and coalescent-based simulation approaches to determine two parameters that are often difficult to infer, particularly for ancestral lineages: the timing and geographic extent of gene flow.

### Phylogeography of the *S. spinosus* Group

A number of phylogeographic studies have been performed in Mexico due to its rich orogenic history (e.g., Devitt 2006; Bryson et al 2011a; Bryson Jr et al 2012a; Leaché et al 2013), and many studies have found that the major mountain ranges (Sierra Madre Occidental, western Mexico; Sierra Madre Oriental, eastern Mexico; Trans-Mexican Volcanic Belt, southern-central Mexico; Sierra Madre del Sur, southern Mexico) have had major effects on the biogeographic patterns across many taxonomic groups (e.g., Bryson Jr et al 2012b; Ruiz-Sanchez & Specht 2013). On the other hand, some researchers argue that some of these features do not represent single biogeographic entities (e.g., Corona et al 2007). Although the extant distribution of *S. spinosus* group taxa is similar to other species (e.g., *Phrynosoma orbiculare*; Bryson Jr et al 2012), subtleties in habitat (and therefore elevational) preferences result in a unique phylogeographic distribution across Mexico for this group, particularly in the geographic overlap of distinct populations in southeastern Mexico (but see Fernández 2011).

Population assignment and species delimitation analyses identified six independent lineages within the *S. spinosus* group (Fig. 3; Table 2); geographic distributions largely coincide with the ranges of subspecies (Figs. 1,3). The geographic boundaries of these lineages appear to be strongly influenced by the geology of the region. In southwestern Mexico, the Rio Santiago, Rio Ahuijullo, and the western portion of the Balsas basins form the interface between northern and central *S. horridus* populations (Figs. 1,3). These barriers have also been implicated in lineage divergence of horned lizards (*Phrynosoma*; Bryson Jr et al 2012b) and rattlesnakes (*Crotalus*; Bryson et al 2011a). We performed a genetic landscape interpolation in the program Alleles in Space to inspect the concordance between geography and the genetic landscape, and the coincidence between these two landscapes was moderately strong (Supplemental Figs. 6,7). The Trans-Mexican Volcanic Belt corresponds to the north-south barrier separating northern and central *S. spinosus* populations (Fig. 1). That this geologic feature is a natural barrier causing population differentiation is no surprise, as the average elevation is 2,300m and many peaks in this range are >3000m (some are >5000m) and habitats are widely varied (Marshall & Liebherr 2000). The low elevation valleys between the Trans-Mexican Volcanic Belt and Sierra Madre del Sur in northwestern Oaxaca and eastern Puebla likewise seem to be isolating southern populations of *S. spinosus*, a pattern seen in other lizard species (Bryson & Riddle 2012).

The time-calibrated species tree indicated that the common ancestor of *S. spinosus* and *S. horridus* diverged approximately 3.1 mya (Fig. 6). This is in agreement with Cole’s (1970) hypothesis that these two species originated in the late Pliocene. Since this time, Mexico has gone through a number of glacial and pluvial (precipitation) cycles causing range expansions and contractions and population coalescence and divergence of many species (Hewitt 2004). Ancestral *S. spinosus* and *S. horridus* populations were isolated to the Central Mexican Plateau and western slope of the Sierra Madre Occidental, respectively, likely due to Pleistocene glacial cycles (Riddle & Hafner 2006). Following separation, pluvial climates allowed the northern and central populations of *S. horridus* to be “in more-or-less continuous contact with each other” (Frost 1978).

The prolonged extent of geographic overlap between ancestral lineages of *S. horridus* and *S. spinosus* provided ample opportunity for genetic exchange between these lineages. However, our simulation results showed that little to no ancestral gene flow occurred in this region (for two out of three lineages modeled; Table 3), which refutes the model of divergence with gene flow. The lack of ancestral gene flow, in spite of our phylogeographic results, could be for a few reasons. First, the ancestral locations of these lineages was incorrectly reconstructed. The method of species tree geographic diffusion is new (Nylinder et al 2014) and has not been tested under simulation, and we are therefore unaware of any inaccuracies it may have. Furthermore, the ancestral locations the method is allowed to explore are limited to the geographic extent of extant distributions (or however else the researcher chooses to draw the population-delimiting polygons prior to analysis). Simulations and further empirical studies must be performed with this method to determine its accuracy. Secondly, individuals within the reconstructed ancestral ranges may have been occupying the (small) regions allopatric/parapatric to the other species. This is possible, however, not likely, as the regions in allopatry are peripheral and small in comparison with each lineages’ entire range. Third, although ancestral *S. spinosus* and *S. horridus* may have been broadly sympatric, they may have not been syntopic. Both species currently inhabit mostly xeric habitats, but show different microhabitat preferences (Cole 1970), meaning they simply may have not historically come into contact. And lastly, perhaps species-specific recognition cues were more pronounced due to reinforcement as ancestral populations diverged. Frost (1978) noted a northwest-southeast cline in *S. h. albiventris/S. h. oligoporus* populations for some external morphological characters (e.g., color patterning) that he posited was due to reinforcement at the subspecific boundary. Such a situation could be a strong barrier to ancestral gene flow.

The phylogeographic model of 2^°^ contact is the most likely given our results, in concert. The simulation modeling 2^°^ contact (Scenario E; Fig. 2) fit the empirical data for southern populations of both *S. spinosus* and *S. horridus*, although the empirical data for southern *S. spinosus* (population 3) did not fit the results from this scenario. Only one “*S. spinosus* south” individual was recovered in the admixed mtDNA clade, potentially indicating a low level of gene flow that did not match the simulations modeling a higher migration rate for this taxon. The split of the common ancestor of southern *S. spinosus* populations into its daughter lineages did not occur until around 860,000 years ago. After this point, the ranges of southern *S. horridus* and *S. spinosus* shared a moderate amount of range overlap in southern Mexico where much of the admixed mtDNA clade is situated (Figs. 4,6). The patterns of, or lack thereof, nDNA ancestral gene flow detected in the IMa2 analyses further support the 2^°^ contact model. No ancestral gene flow was detected between *S. spinosus* and *S. horridus* common ancestors, but was detected between extant populations of *S. spinosus* and *S. horridus* (Table 4). Although a new study by Leaché et al (2013a) found evidence for divergence with gene flow between *S. horridus* and *S. spinosus*, the method they used did not allow for discernment between models of 2^°^ vs. divergence with gene flow.

### Mito-nuclear Discordance in the *S. spinosus* Group

Numerous studies have reported conflicting evolutionary histories between nuclear and mitochondrial genomes (“mito-nuclear discordance”, reviewed in Toews & Brelsford 2012). Out of 126 studies identified by Toews & Brelsford (2012) that documented strong incongruence between mt- and nDNA biogeographic patterns, the overwhelming majority of cases (97%) reported that the discordance likely arose from geographic isolation followed by secondary contact; the most common form of mito-nuclear discordance is due to the asymmetric movement of mtDNA between lineages. In the case of the *S. spinosus* group, we can safely rule out the possibility that incomplete lineage sorting (ILS) of mtDNA alleles as the cause of mito-nuclear incongruence, as we would expect ILS to leave a geographically-independent genealogical signature. We cannot, however, rule out the possibility that adaptive introgression may be a factor, particularly because many of the individuals belonging to the “admixed” mtDNA clade were collected in moderately high elevation sites (>2000m) where individuals with particular mitochondrial haplotypes may be better adapted Cheviron and Brumfield, 2009.

The most likely cause of mito-nuclear discordance in the *S. spinosus* group appears to be due to unidirectional gene flow from southern *S. spinosus* and *S. horridus* into central *S. spinosus*. The admixed mtDNA clade is composed of central *S. spinosus*, southern *S. spinosus*, and southern *S. horridus* individuals (Figs. 1,4). Whereas southern *S. horridus* and *S. spinosus* individuals were recovered in other mitochondrial clades, all central *S. spinosus* individuals were confined to the admixed clade. This phylogenetic pattern suggests that the admixed mtDNA clade was originally composed of all central *S. spinosus* individuals, and recently, that southern *S. horridus* and *S. spinosus* males have introgressed their mtDNA copies into central *S. spinosus* females. Our gsi results support the notion of recent (mitochondrial) gene flow between southern *S. horridus* and central *S. spinosus*, but not between southern *S. spinosus* and central *S. spinosus* (Table 3).

### Population Distinctiveness and Divergence Within the *Sceloporus spinosus* Group

Distinguishing between what have been considered distinct taxa in the *S. spinosus* group is problematic, and little agreement exists between previous authors. For instance, Boyer et al (1987) concluded that *S. s. spinosus* and *S. h. horridus* were conspecific based on the overlap of femoral pores and contact frequency of supraocular-median head scales as a result of intergradation. Smith & Chiszar (1992) later returned *S. spinosus* and *S. horridus* to specific status after a reinterpretation of these individuals as intergrades between *S. s. spinosus* and *S. s. apicalis.* Distinguishing between *S. spinosus* subspecies is also problematic due to the slight difference in average values of quantitative characters between subspecies, where Smith & Chiszar (1992) note that examination only of a series of six or more permits ”reasonably secure identifications”. Similar problems exist within *S. horridus*, where Frost (1978) reported a large area of intergrade in western Mexico between *S. h.* albiventris and *S. h. oligoporus*. Notwithstanding, Lemos-Espinal et al (2004) regarded these taxa as distinct species.

Contrary to some previous research (Wiens & Reeder 1997; Smith 2001), the results of our nDNA-based phylogeny show the *S. spinosus* group to be monophyletic (to the exclusion of *S. edwardtaylori*; Fig. 4). Furthermore, *S. spinosus* and *S. horridus* are monophyletic with respect to each other, a result at odds with previous research (Wiens et al 2010). This discrepancy is certainly due to the overriding signal of the mtDNA in the combined mt- and nDNA analysis of Wiens et al (2010). Lineages are often determined to be distinct based on an assessment of gene flow levels, a test of the biological species concept (Mayr 1942; Mayr et al 1963). Our tests of nuclear gene flow in the *S. spinosus* group revealed gene flow not only between populations of each species, but also across species (Table 4). But, the level of gene flow we detected in all instances was far below 0.5 *N*_*e*_*m* per generation, a value used by some when determining species limits (Porter 1990; Hey 2009). The interpretation of these results, particularly for the 6-population model, should be cautioned because the size of our molecular dataset is likely inadequate to generate accurate results (Hey 2010; Choi & Hey 2011).

We based our species delimitation models on a combination of results from population assignment and migration analyses. In an attempt to account for gene flow, which has been show to severely affect parameter estimation in coalescent-based species tree analyses (Leaché 2009; Leaché et al 2013b), we excluded individuals located near population boundaries (Fig. 3). This of course assumes that the gene flow we detected occurred on population boundaries, an assumption which may not be true. Removing these peripheral individuals did not affect our species delimitation results that indicated the presence of six independent lineages in the *S. spinosus* group.

When comparing gsi values from our coalescent simulations against our empirical mtDNA, it appears that the empirical data for the southern *S. horridus* population are most in agreement with scenarios modeling recent, but not ancestral, gene flow across species (Scenarios C-E; Fig. 2; Table 3). On the other hand, the empirical data of the southernmost *S. spinosus* population are in agreement with a scenario in which there was either no gene flow, or ancestral gene flow between *S. spinosus* and *S. horridus* ancestors. The gsi simulation results did not reject any scenario for the central *S. spinosus* population (Table 3). Our conclusions based on the gsi results are directly a function of the levels of gene flow used in our simulations. We used a relatively high migration value of 1 *N*_*e*_*m* in our simulations (0.5 *N*_*e*_*m* unidirectionally from each population), where some researchers consider a migration rate of *N*_*e*_*m* >0.5 enough to keep populations from diverging (Porter 1990). We therefore believe that we have modeled a realistic level of gene flow to assess matrilineal-based migration in the *S. spinosus* group.

### Conclusions

The number of plausible models that should be evaluated in phylogeographic studies is nearly infinite (e.g., Tsai & Carstens 2013; Pelletier & Carstens 2014). Here, we generated a small number of plausible models based on the results from our empirical data. Given our results, we conclude that i) six independent genetic lineages exist in the *S. spinosus* group, and identifying species is important for accurately modeling evolutionary histories, ii) coalescent simulations reject a model of ancestral gene flow in the *S. spinosus* group, iii) the Bayesian phylogeographic reconstruction for the ancestral ranges of the *S. spinosus* group suggests that species within the group broadly overlapped throughout a majority of their evolutionary history (~3 million years), and iv) mitochondrial introgression is localized spatially, and likely temporally as well. The contrasting evolutionary histories of the nuclear and mitochondrial genomes seem to indicate another example of the mtDNA locus not accurately representing the true species-level evolutionary history. However, the mitochondrial genome has nonetheless provided a valuable piece of information in determining the evolutionary history of the *S. spinosus* group by presenting evidence for the timing and geographic extent of contact between distinct populations in this group.

## Acknowledgments

We’d like to personally thank Dr. Philippe Lemey for his assistance with our Bayesian phytogeography and SPREAD analyses. Similarly, we thank Adam Bazinet for his assistance in our R implementation of the genealogical sorting index package and calculations. This work was facilitated through the use of advanced computational, storage, and networking infrastructure provided by the Hyak supercomputer system at the University of Washington through grant support from the National Science Foundation (DEB-1144630) to ADL. MC was supported by a DGEP PhD and DGAPA postdoctoral scholarships from the Universidad Nacional Autónoma de México (UNAM). Financial support was provided by a grant from the UNAM to FMC (PAPIIT IN2009-01) and the American Museum of Natural History through the Theodore Roosevelt Memorial Grants program. Further support for this project was provided by grants from DGAPA, UNAM (PAPIIT no. IN224009) and CONACYT (no. 154093) to A. Nieto-Montes de Oca. We thank the Museo de Zoología Alfonso L. Herrera (MZFC) and the Colección Nacional de Anfibios y Reptiles (CNAR), UNAM, for the loan of specimens and to MZFC for providing tissue samples. Norma Manríquez and Laura Márquez provided useful advice on lab work. J. C. Barajas, E. Pérez Ramos, R. Meza, G. Zamora, A. Ortega, S. López, V. Serrano, N. Martínez, C. Peña and G. Barrios assisted with field work.

## Data Accessibility

- DNA Sequences: Genbank accession nos. xxxx-xxxx
- Bayesian phylogeography species tree .xml provided at Dryad doi: 10.5061/dryad.3v55p
- All collecting locality information is available in the Supplementary Materials section.

## Author Contributions

All authors designed the research and collected specimens; JAG and MLC obtained the data and conducted analyses; JAG wrote the paper, and all co-authors contributed to editing the manuscript.

## References

Abbott R, Albach D, Ansell S, et al (2013) Hybridization and speciation. Journal of Evolutionary Biology, 26, 229–246.

Alfaro ME, Zoller S, Lutzoni F (2003) Bayes or bootstrap? a simulation study comparing the performance of bayesian markov chain monte carlo sampling and bootstrapping in assessing phylogenetic confidence. Molecular Biology and Evolution, 20, 255–266.

Arèvalo E, Davis SK, Sites JW (1994) Mitochondrial dna sequence divergence and phylogenetic relationships among eight chromosome races of the *Sceloporus grammicus* complex (phrynosomatidae) in central mexico. Systematic Biology, 43, 387–418.

Avise JC (2000) Phylogeography: the history and formation of species. Harvard University Press.

Baele G, Lemey P, Bedford T, Rambaut A, Suchard MA, Alekseyenko AV (2012) Improving the accuracy of demographic and molecular clock model comparison while accommodating phylogenetic uncertainty. Molecular biology and evolution, 29, 2157–2167.

Barley AJ, White J, Diesmos AC, Brown RM (2013) The challenge of species delimitation at the extremes: diversification without morphological change in philippine sun skinks. Evolution, 67, 3556–3572.

Bielejec F, Rambaut A, Suchard MA, Lemey P (2011) Spread: Spatial phylogenetic reconstruction of evolutionary dynamics. Bioinformatics.

Boyer T, Smith H, Casas-Andreu G (1987) The taxonomic relationships of the mexican lizard species *Sceloporus horridus*. Bulletin of the Maryland Herpetological Society, 18, 189–191.

Brandley MC, Wang Y, Guo X, et al (2011) Accommodating heterogenous rates of evolution in molecular divergence dating methods: an example using intercontinental dispersal of plestiodon (eumeces) lizards. Systematic Biology, 60, 3–15.

Bryson RW, García-Vázquez UO, Riddle BR (2011a) Phylogeography of middle american gophersnakes: mixed responses to biogeographical barriers across the mexican transition zone. Journal of Biogeography, 38, 1570–1584.

Bryson RW, Murphy RW, Lathrop A, Lazcano-Villareal D (2011b) Evolutionary drivers of phylogeographical diversity in the highlands of mexico: a case study of the *Crotalus triseriatus* species group of montane rattlesnakes. Journal of Biogeography, 38, 697–710.

Bryson RW, Riddle BR (2012) Tracing the origins of widespread highland species: a case of neogene diversification across the mexican sierras in an endemic lizard. Biological Journal of the Linnean Society, 105, 382–394.

Bryson Jr RW, García-Vázquez UO, Riddle BR (2012a) Diversification in the mexican horned lizard *Phrynosoma orbiculare* across a dynamic landscape. Molecular phylogenetics and evolution, 62, 87–96.

Bryson Jr RW, García-Vázquez UO, Riddle BR (2012b) Relative roles of neogene vicariance and quaternary climate change on the historical diversification of bunchgrass lizards (*Sceloporus scalaris* group) in mexico. Molecular phylogenetics and evolution, 62, 447–457.

Carstens B, Degenhardt J, Stevenson A, Sullivan J (2005) Accounting for coalescent stochasticity in testing phylogeographical hypotheses: modelling pleistocene population structure in the idaho giant salamander dicamptodon aterrimus. Molecular Ecology, 14, 255–265.

Chan LM, Brown JL, Yoder AD (2011) Integrating statistical genetic and geospatial methods brings new power to phylogeography. Molecular phylogenetics and evolution, 59, 523–537.

Chiari Y, van der Meijden A, Mucedda M, Lourenço JM, Hochkirch A, Veith M (2012) Phylogeography of sardinian cave salamanders (genus *Hydromantes*) is mainly determined by geomorphology. PloS one, 7, e32332.

Choi SC, Hey J (2011) Joint inference of population assignment and demographic history. Genetics, 189, 561–577.

Cole CJ (1970) Karyotypes and evolution of the *spinosus* group of lizards in the genus *Sceloporus*. American Museum Novitates, 2431, 1–47.

Corona AM, Toledo VH, Morrone JJ (2007) Does the trans-mexican volcanic belt represent a natural biogeographical unit? an analysis of the distributional patterns of coleoptera. Journal of Biogeography, 34, 1008–1015.

Cummings MP, Neel MC, Shaw KL (2008) A genealogical approach to quantifying lineage divergence. Evolution, 62, 2411–2422.

Darriba D, Taboada GL, Doallo R, Posada D (2012) jmodeltest 2: more models, new heuristics and parallel computing. Nature Methods, 9, 772–772.

Darwin C (1859) The origin of species by means of natural selection: or, the preservation of favored races in the struggle for life. John Murray.

Devitt TJ (2006) Phylogeography of the western lyresnake (trimorphodon biscutatus): testing aridland biogeographical hypotheses across the nearctic–neotropical transition. Molecular Ecology, 15, 4387–4407.

Drummond AJ, Rambaut A (2007) Beast: Bayesian evolutionary analysis by sampling trees. BMC evolutionary biology, 7, 214.

Excoffier L, Lischer HE (2010) Arlequin suite ver 3.5: a new series of programs to perform population genetics analyses under linux and windows. Molecular ecology resources, 10, 564–567.

Fan Y, Wu R, Chen MH, Kuo L, Lewis PO (2011) Choosing among partition models in bayesian phylogenetics. Molecular biology and evolution, 28, 523–532.

Fernández JA (2011) Comparative biogeography of the arid lands of Central Mexico. Ph.D. thesis, Louisiana State University.

Frost D (1978) Geographic variation and evolution of Sceloporus horridus Wiegmann (Lacertilia: Iguanidae). Master’s thesis, Louisiana State University.

Grummer JA, Bryson Jr RW, Reeder TW (2014) Species delimitation using bayes factors: simulations and application to the *Sceloporus scalaris* species group (squamata: Phrynosomatidae). Systematic Biology, 63, 119–133.

Guillot G (2008) Inference of structure in subdivided populations at low levels of genetic differentiation—the correlated allele frequencies model revisited. Bioinformatics, 24, 2222–2228.

Guillot G, Estoup A, Mortier F, Cosson JF (2005a) A spatial statistical model for landscape genetics. Genetics, 170, 1261–1280.

Guillot G, Mortier F, Estoup A (2005b) Geneland: a computer package for landscape genetics. Molecular Ecology Notes, 5, 712–715.

Guindon S, Gascuel O (2003) A simple, fast, and accurate algorithm to estimate large phylogenies by maximum likelihood. Systematic biology, 52, 696–704.

Harrison RG (1993) Hybrid zones and the evolutionary process. Oxford University Press.

Heled J, Drummond AJ (2010) Bayesian inference of species trees from multilocus data. Molecular biology and evolution, 27, 570–580.

Hewitt G (2004) Genetic consequences of climatic oscillations in the quaternary. Philosophical Transactions of the Royal Society of London. Series B: Biological Sciences, 359, 183–195.

Hewitt GM (2001) Speciation, hybrid zones and phylogeography—or seeing genes in space and time. Molecular Ecology, 10, 537–549.

Hey J (2009) On the arbitrary identification of real species. Speciation and Patterns of Diversity (eds Butlin RK, Bridle J, Schluter D), pp. 15–28.

Hey J (2010) Isolation with migration models for more than two populations. Molecular biology and evolution, 27, 905–920.

Hickerson M, Carstens B, Cavender-Bares J, et al (2010) Phylogeography‘s past, present, and future: 10 years after. Molecular Phylogenetics and Evolution, 54, 291–301.

Hillis DM, Bull JJ (1993) An empirical test of bootstrapping as a method for assessing confidence in phylogenetic analysis. Systematic biology, 42, 182–192.

Hudson RR (2002) Generating samples under a wright–fisher neutral model of genetic variation. Bioinformatics, 18, 337–338.

Huelsenbeck JP, Andolfatto P (2007) Inference of population structure under a dirichlet process model. Genetics, 175, 1787–1802.

Huelsenbeck JP, Andolfatto P, Huelsenbeck ET (2011) Structurama: Bayesian inference of population structure. Evolutionary bioinformatics online, 7, 55.

Jakobsson M, Rosenberg NA (2007) Clumpp: a cluster matching and permutation program for dealing with label switching and multimodality in analysis of population structure. Bioinformatics, 23, 1801–1806.

Kass RE, Raftery AE (1995) Bayes factors. Journal of the american statistical association, 90, 773–795.

Knowles LL (2001) Did the pleistocene glaciations promote divergence? tests of explicit refugial models in montane grasshopprers. Molecular Ecology, 10, 691–701.

Knowles LL (2009) Statistical phylogeography. Annual Review of Ecology, Evolution, and Systematics, 40, 593–612.

Kocher TD, Thomas WK, Meyer A, et al (1989) Dynamics of mitochondrial dna evolution in animals: amplification and sequencing with conserved primers. Proceedings of the National Academy of Sciences, 86, 6196–6200.

Kuhner MK (2009) Coalescent genealogy samplers: windows into population history. Trends in Ecology & Evolution, 24, 86–93.

Larson EL, White TA, Ross CL, Harrison RG (2013) Gene flow and the maintenance of species boundaries. Molecular Ecology.

Leaché A, Sites Jr J (2010) Chromosome evolution and diversification in north american spiny lizards (genus sceloporus). Cytogenetic and genome research, 127, 166–181.

Leaché AD (2009) Species tree discordance traces to phylogeographic clade boundaries in north american fence lizards (sceloporus). Systematic Biology, p. syp057.

Leaché AD (2010) Species trees for spiny lizards (genus *Sceloporus*): Identifying points of concordance and conflict between nuclear and mitochondrial data. Molecular phylogenetics and evolution, 54, 162–171.

Leaché AD Harris RB, Maliska ME, Linkem CW (2013a) Comparative species divergence across eight triplets of spiny lizards (sceloporus) using genomic sequence data. Genome biology and evolution, 5, 2410–2419.

Leaché AD Harris RB, Rannala B, Yang Z (2013b) The influence of gene flow on species tree estimation: a simulation study. Systematic biology, p. syt049.

Leaché AD Palacios JA, Minin VN, Bryson RW (2013) Phylogeography of the trans-volcanic bunchgrass lizard (*Sceloporus bicanthalis*) across the highlands of south-eastern mexico. Biological Journal of the Linnean Society, 110, 852–865.

Leaché AD Reeder TW (2002) Molecular systematics of the eastern fence lizard (sceloporus undulatus): a comparison of parsimony, likelihood, and bayesian approaches. Systematic biology, 51, 44–68.

Lemey P, Rambaut A, Drummond AJ, Suchard MA (2009) Bayesian phylogeography finds its roots. PLoS computational biology, 5, e1000520.

Lemey P, Rambaut A, Welch JJ, Suchard MA (2010) Phylogeography takes a relaxed random walk in continuous space and time. Molecular biology and evolution, 27, 1877–1885.

Lemos-Espinal J, Chiszar D, Smith H, Woolrich-Piña G (2004) Selected records of 2003: lizards from chihuahua and sonora, mexico. Bulletin of the Chicago Herpetological Society, 39, 164–168.

Maddison W, Maddison D (2011) Mesquite 2.75: a modular system for evolutionary analysis.

Mallet J (2005) Hybridization as an invasion of the genome. Trends in Ecology & Evolution, 20, 229–237.

Marshall C, Liebherr J (2000) Cladistic biogeography of the mexican transition zone. Journal of biogeography, 27, 203–216.

Mayr E (1942) Systematics and the origin of species, from the viewpoint of a zoologist. Harvard University Press.

Mayr E, et al (1963) Animal species and evolution. Animal species and their evolution.

McGuire G, Wright F (2000) Topal 2.0: improved detection of mosaic sequences within multiple alignments. Bioinformatics, 16, 130–134.

Miller M (2005) Alleles in space (ais): computer software for the joint analysis of interindividual spatial and genetic information. Journal of Heredity, 96, 722–724.

Milne I, Lindner D, Bayer M, et al (2009) Topali v2: a rich graphical interface for evolutionary analyses of multiple alignments on hpc clusters and multi-core desktops. Bioinformatics, 25, 126–127.

Nielsen R, Wakeley J (2001) Distinguishing migration from isolation: a markov chain monte carlo approach. Genetics, 158, 885–896.

Nylinder S, Lemey P, De Bruyn M, et al (2014) On the biogeography of centipeda: A species-tree diffusion approach. Systematic biology, 63, 178–191.

Pelletier TA, Carstens BC (2014) Model choice for phylogeographic inference using a large set of models. Molecular ecology.

Pettengill JB, Moeller DA (2012) Phylogeography of speciation: allopatric divergence and secondary contact between outcrossing and selfing *Clarkia*. Molecular Ecology, 21, 4578–4592.

Porter AH (1990) Testing nominal species boundaries using gene flow statistics: the taxonomy of two hybridizing admiral butterflies (limenitis: Nymphalidae). Systematic Biology, 39, 131–147.

Portik DM, Wood PL Jr, Grismer JL, Stanley EL, Jackman TR (2012) Identification of 104 rapidly-evolving nuclear protein-coding markers for amplification across scaled reptiles using genomic resources. Conservation Genetics Resources, 4, 1–10.

Rambaut A, Drummond A (2007) Tracer v1. 4.

Rannala B, Yang Z (2003) Bayes estimation of species divergence times and ancestral population sizes using dna sequences from multiple loci. Genetics, 164, 1645–1656.

Riddle B, Hafner D (2006) A step-wise approach to integrating phylogeographic and phylogenetic biogeographic perspectives on the history of a core north american warm deserts biota. Journal of Arid Environments, 66, 435–461.

Ronquist F, Huelsenbeck JP (2003) Mrbayes 3: Bayesian phylogenetic inference under mixed models. Bioinformatics, 19, 1572–1574.

Rosenblum E, Belfiore N, Moritz C (2007) Anonymous nuclear marker discovery in the eastern fence lizard, *Sceloporus undulatus*. Molecular Ecology Notes, 7, 113–116.

Ruiz-Sanchez E, Specht CD (2013) Influence of the geological history of the trans-mexican volcanic belt on the diversification of nolina parviflora (asparagaceae: Nolinoideae). Journal of Biogeography, 40, 1336–1347.

Smith CI, Tank S, Godsoe W, et al (2011) Comparative phylogeography of a coevolved community: concerted population expansions in joshua trees and four *Yucca* moths. PloS one, 6, e25628.

Smith E (2001) Species boundaries and evolutionary patterns of speciation among the malachite lizards formosus group of the genus Sceloporus (Squamata: Phrynosomatidae). Ph.D. thesis, University of Texas at Arlington.

Smith H (1939) The Mexican and Central American Lizards of the Genus *Sceloporus*, vol. 26 of Zoological Series. Field Museum Press.

Smith H, Chiszar D (1992) *Sceloporus horridus* and *S. spinosus* (reptilia: Sauria) are separate species. Bulletin of the Maryland Herpetological Society, 2, 44–52.

Smith P, Smith H (1951) A new lizard (Sceloporus) from oaxaca, mexico. Proceedings of the Biological Society of Washington, 64, 101–104.

Spellman GM, Klicka J (2006) Testing hypotheses of pleistocene population history using coalescent simulations: phylogeography of the pygmy nuthatch (*Sitta pygmaea*). Proceedings of the Royal Society B: Biological Sciences, 273, 3057–3063.

Stamatakis A (2006) Raxml-vi-hpc: maximum likelihood-based phylogenetic analyses with thousands of taxa and mixed models. Bioinformatics, 22, 2688–2690.

Stephens M, Smith NJ, Donnelly P (2001) A new statistical method for haplotype reconstruction from population data. The American Journal of Human Genetics, 68, 978–989.

Toews DP, Brelsford A (2012) The biogeography of mitochondrial and nuclear discordance in animals. Molecular Ecology, 21, 3907–3930.

Townsend TM, Alegre RE, Kelley ST, Wiens JJ, Reeder TW (2008) Rapid development of multiple nuclear loci for phylogenetic analysis using genomic resources: an example from squamate reptiles. Molecular phylogenetics and evolution, 47, 129–142.

Tsai YHE, Carstens BC (2013) Assessing model fit in phylogeographical investigations: an example from the north american sandbar willow salix melanopsis. Journal of Biogeography, 40, 131–141.

Wiens JJ, Kuczynski CA, Arif S, Reeder TW (2010) Phylogenetic relationships of phrynosomatid lizards based on nuclear and mitochondrial data, and a revised phylogeny for *Sceloporus*. Molecular Phylogenetics and Evolution, 54, 150–161.

Wiens JJ, Reeder TW (1997) Phylogeny of the spiny lizards (*Sceloporus*) based on molecular and morphological evidence. Herpetological Monographs, pp. 1–101.

Xie W, Lewis PO, Fan Y, Kuo L, Chen MH (2011) Improving marginal likelihood estimation for bayesian phylogenetic model selection. Systematic Biology, 60, 150–160.

Yang Z, Rannala B (2010) Bayesian species delimitation using multilocus sequence data. Proceedings of the National Academy of Sciences, 107, 9264–9269.

